# Modernizing histopathological analysis: a fully automated workflow for the digital image analysis of the intestinal microcolony survival assay

**DOI:** 10.1101/2024.12.09.627578

**Authors:** Alexander Baikalov, Ethan Wang, Denae Neill, Nihar Shetty, Trey Waldrop, Kevin Liu, Abagail Delahousessaye, Edgardo Aguilar, Nefetiti Mims, Stefan Bartzsch, Emil Schüler

## Abstract

**Background:** Manual analysis of histopathological images is often not only time-consuming and painstaking but also prone to error from subjective evaluation criteria and human error. To address these issues, we created a fully automated workflow to enumerate jejunal crypts in a microcolony survival assay to quantify gastrointestinal damage from radiation.

**Methods and Materials:** After abdominal irradiation of mice, jejuna were obtained and prepared on histopathologic slides, and crypts were counted manually by trained individuals. The automated workflow (AW) involved obtaining images of jejunal slices from the irradiated mice, followed by cropping and normalizing the individual slice images for resolution and color; using deep learning-based semantic image segmentation to detect crypts on each slice; using a tailored algorithm to enumerate the crypts; and tabulating and saving the results. A graphical user interface (GUI) was developed to allow users to review and correct the automated results.

**Results:** Crypts counted manually exhibited a mean absolute percent deviation of (34 ± 26)% between individuals vs the group mean across counters, which was reduced to (11 ± 6)% across the 3 most-experienced counters. The AW processed a sample image dataset from 60 mice in a few hours and required only a few minutes of active user effort. AW counts deviated from experts’ mean counts by (10 ± 8)%. The AW thereby allowed rapid, automated evaluation of the microcolony survival assay with accuracy comparable to that of trained experts and without subjective inter-observer variation.

**Highlights:**

- We fully automated the digital image analysis of a microcolony survival assay
- Analyzing 540 images takes a few hours with only minutes of active user effort
- The automated workflow (AW) is just as accurate as trained experts
- The AW eliminates subjective inter-observer variation and human error
- Human review possible with built-in graphical user interface

**Graphical Abstract:** 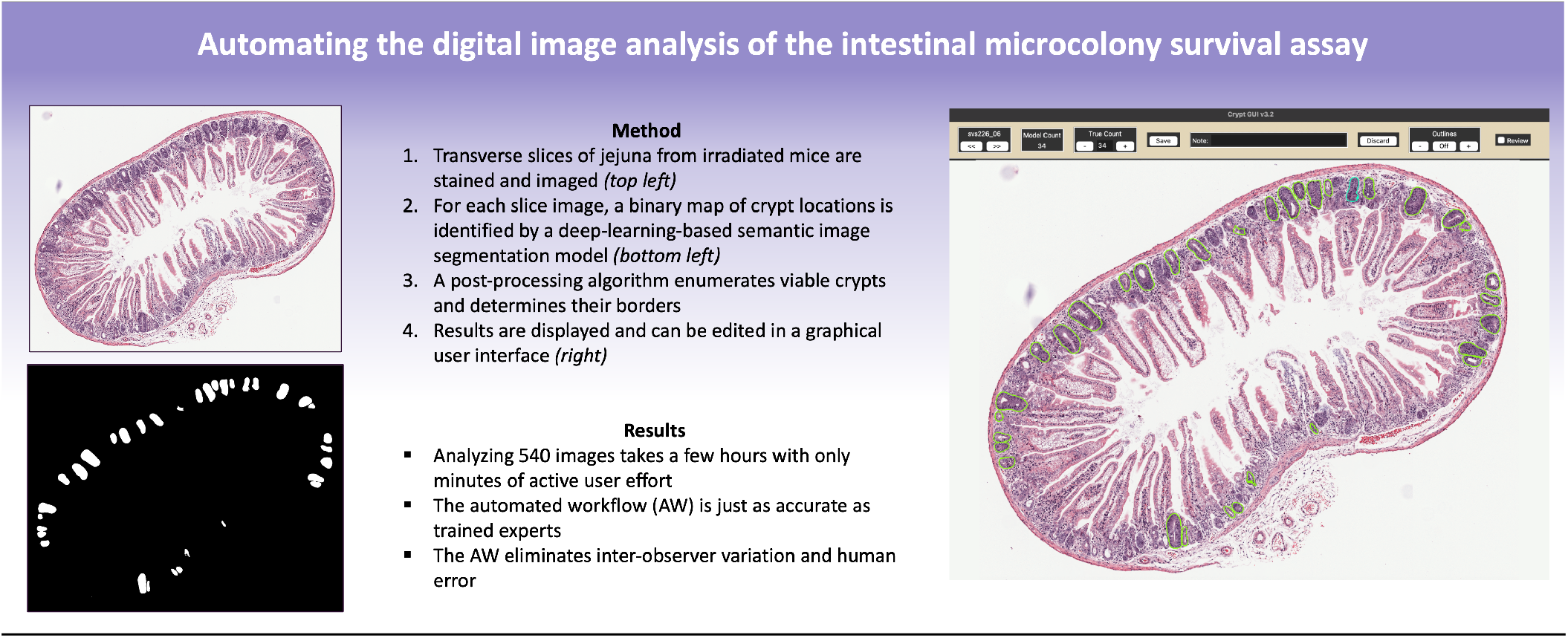

## Introduction

In radiation biology research, a gold standard assay for evaluating radiation-induced gastrointestinal toxicity in mice is the microcolony survival assay, which equates the number of regenerating jejunal crypts to the extent of radiation damage [1]. Since its establishment in 1970, the microcolony survival assay has been preferred for its direct, in-vivo assessment of cellular toxicity and rapid turnaround time relative to clonogenic assays [2]. It also allows assessments of the effect of potential radio-modulatory factors such as oxygenation [1] and chemotherapy [3-6] on radiation response via comparisons of dose-response curves. However, regenerating crypts are generally quantified manually, which is time-consuming, subjective, and inevitably introduces variability in the analyzed data set, as has been noted for other manual quantification assays [7]. Moreover, manual analysis may limit the amount of data that can be reasonably analyzed. “Representative data sets” are often used instead, which introduces the risk of missing, understating, or overstating key findings in the analyzed data set [3].

The advent of digital whole-slide imaging, in which sequential images of glass slides are ‘stitched’ together to form virtual whole-slide images (WSIs) [8], has expanded the possibilities of histologic slide analysis, particularly the use of artificial intelligence (AI) algorithms to automate data extraction from WSIs [9, 10].

Automation of jejunal crypt counting is highly desirable for facilitating consistent crypt counting, reducing manual labor, and allowing analyses of WSIs in their entirety. A previous study showcased a deep-learning model trained to estimate radiation dose based on WSI of jejunal slices [11] that provided an automated alternative to the typical microcolony survival assay for radiation damage. However, the features indicating greater or lesser dose burden as determined by the model are unknown and may not correlate with biological features. Although such a model may be useful in specific applications, it cannot replace the standard microcolony survival assay.

In this study, we demonstrate a novel, AI-based framework to fully automate the microcolony survival assay based on WSIs of mouse jejunal slices. Our framework consists of four steps: (1) Digital image preparation, (2) Image segmentation, (3) Feature quantification, and (4) Data visualization and review. Image segmentation is performed by nnU-Net, a U-Net-based deep learning framework for biomedical image segmentation [12]. nnU-Net was selected because it can be generalized to a variety of segmentation tasks, implemented easily, and has demonstrated success in WSI segmentation [12-14]. Our goals in establishing this automated framework were to functionally replace the manual assay to reduce the time, manual labor, and subjective inconsistency inherent to un-aided manual crypt counting. Achieving this aim would globally facilitate the implementation of this valuable assay by eliminating the need for substantial training and time commitment, as well as improving inter-institutional standardization and thus comparisons of assay findings. Second, we sought to consider data that are traditionally difficult to obtain from physical glass slides (e.g. crypt area, jejunal circumference) and thus are not included in standard microcolony survival assays, and to use these data to better characterize jejunal response to radiation toxicity. Finally, by implementing the nnU-Net framework without task-specific tailoring but applying generic post-processing algorithms, we hope to demonstrate that WSI structure enumeration can be achieved across a variety of clinical and research tasks and require minimal foundational knowledge in AI algorithm construction.

## Methods

### Metrics of data comparison

Throughout this work, we compare crypt counts of jejunal image datasets obtained with different counting methods. To quantify the deviation of a set of crypt counts X from a set of crypt counts Y, we report two metrics: the mean absolute deviation 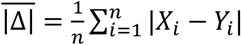 and the mean absolute percent deviation 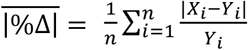. Interpreting 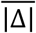 is sometimes difficult owing to the large range of values of crypt counts and thus we included 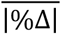. However, the discrete nature of the crypt count metric means that 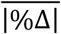is disproportionately skewed high by slices with low (e.g., <10) counts. Therefore, it is instead reported only for slices with counts >10 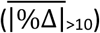.

### Sample preparation, imaging, and manual counting protocols

This work examines microcolony survival assay samples from several datasets as described in **Table 1**. All samples were prepared as follows. Each C57BL/6J mouse was euthanized at 84 ± 4 hours after total abdominal irradiation and its jejunum harvested as follows. An opening was made in the peritoneal cavity, the small intestines were cut at the duodenojejunal junction, and the jejunum was carefully removed with a final cut roughly 8 cm from the first cut. The jejunum was flushed with phosphate-buffered saline to remove all debris, held in place on cardstock, and placed in 10% formalin in a 4°C fridge for 24 hours, after which it was transferred to a 70% ethanol solution. The proximal jejunum was then cut into nine 3-5-mm-long segments **(Fig. 1a)**, which were placed into inserts of a custom 3D-printed cassette. The cassette held the segments upright and next to each other, such that subsequent mounting of the cassette in paraffin yielded a single block containing all nine segments **(Fig. 1b)**. The orientation of the segments in the cassette also preserved their order along the jejunum. A cryostat was used to cut the block of segments into 3-µm-thick transverse sections **(Fig. 1c)**. One section from approximately the center of the block was retained **(Fig. 1d)**, stained with hematoxylin and eosin (H&E), and imaged with an Aperio ScanScope at 20x magnification **(Fig. 1e)**. The ensuing WSI of the section was saved in Aperio ScanScope Virtual Slide (.SVS) file format, resulting in a digital file size of approximately 100–200 MB per WSI and a resolution of 0.5 µm/pixel. This sample preparation protocol thereby yields, for each mouse, one image of nine 3-µm-thick transverse jejunal cross-sections, which we refer to as jejunal slices, evenly spaced along the first 2.7-4.5 cm of the proximal jejunum of the mouse. These slices are numbered 1-9, in increasing order along the length of the jejunum.

**Table 1.** Datasets of jejunal slice images examined and corresonding analyses. Mice were irradiated with a variety of doses and thus exhibited a range of crypt counts. For some mice, only a subset of their 9 jejunal slices were considered. Dataset A2 is a subset of Dataset A as counted by 3 experts. The manual workflow was performed independently for all datasets except for Dataset B, for which the 3 experts performed the manual workflow together and reached a consensus crypt count for each image.

| Dataset | Mice | Jejunal Slices | Range of Crypt Counts | Individuals performing Manual Counting Workflow | Purpose |
| --- | --- | --- | --- | --- | --- |
| **A** | 38 | 342 | 0–120 | 5–10, both experts and non-experts | Quantification of manual workflow variance |
| **A2** | 30 | 270 | 0–120 | 3 experts | Comparison to automated workflow crypt counts |
| **B** | 19 | 59 | 0–83 | 3 experts (consensus) | Training/test datasets for the segmentation model of the automated workflow |
| **C** | 44 | 100 | 0–30 | 1 expert | Human review of automated workflow crypt counts and error type analysis |
| **D** | 34 | 306 | 0–68 | 1 expert | Human review of automated workflow crypt counts and error type analysis |

**Figure 1.**
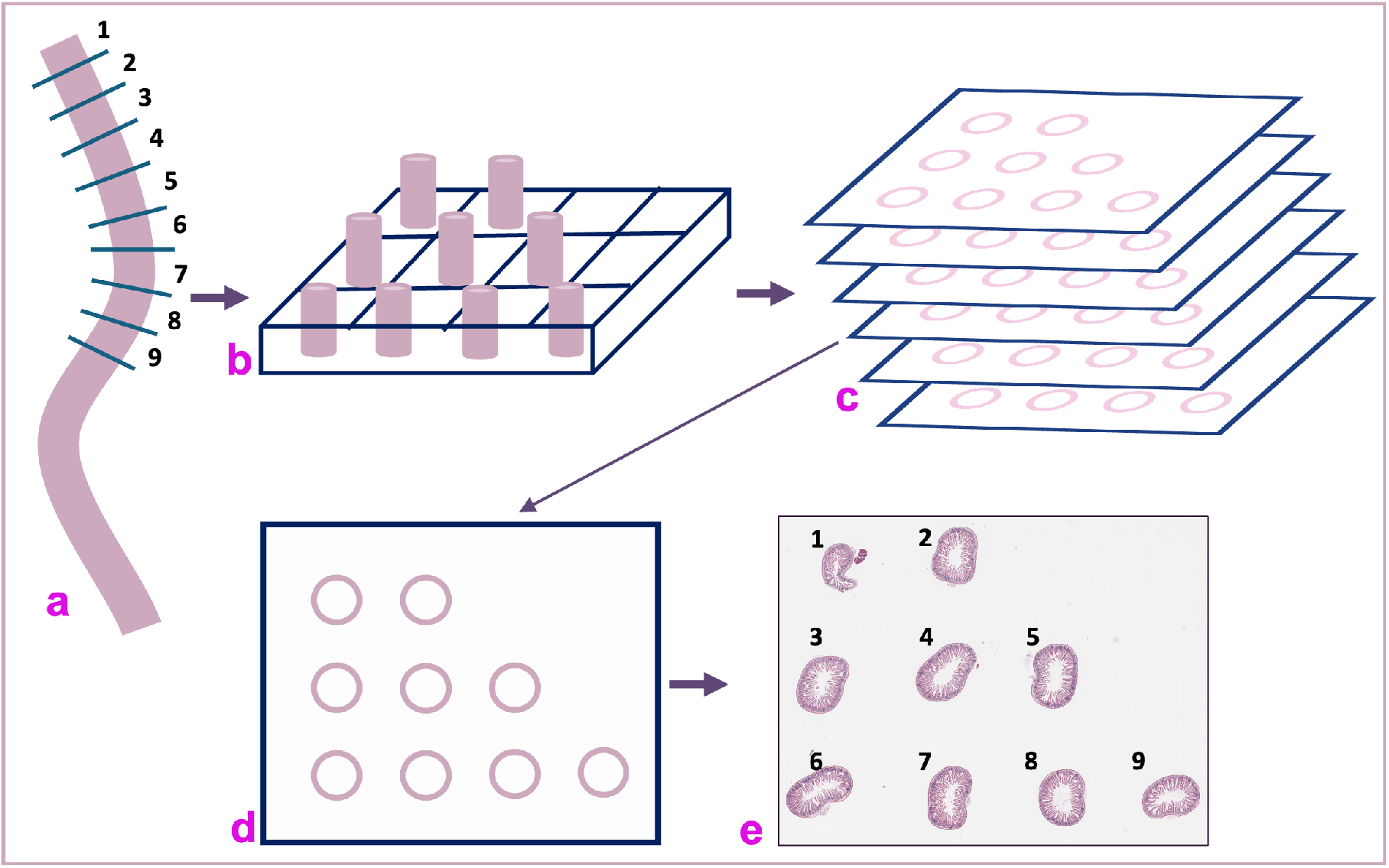
Sample preparation workflow. **(a)** The jejunum of each mouse is cut into 9 segments, which are **(b)** placed upright in a cassette and mounted in paraffin and **(c)** cut into 3-m-thick sections. **(d)** One section from the center of the block is retained, stained, and imaged to yield **(e)** an image of 9 jejunal slices.

The manual crypt counting workflow (MW) was as follows. 3D Slicer software was used to open each WSI and zoom into each of the 9 jejunal slices [15]. If a slice was damaged, incomplete, or improperly imaged, it was dismissed. The individual counter would follow along the circumference of the slice and identify and count crypts. Counts were recorded in an Excel file. The criteria for a crypt to be counted were those originally proposed by Withers and Elkind as follows: ?10 cells, basophilic, clear structure, located on the circumference of the slice (if it was pushed back by a blood vessel or Peyer’s patch, it still counted) [1]. In total, 15 individuals with various levels of experience were trained on the MW. All individuals underwent at least 3 separate training sessions under the supervision of an expert counter. The 3 expert counters had several years of prior experience with the MW.

### Evaluation of the manual crypt counting workflow

To quantify the inter-observer variance of the MW, we analyzed the manually counted number of crypts in Dataset A (see **Table 1**). Each slice of the dataset was separately counted by 5 to 10 individuals. The 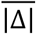and 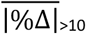 of each individual’s counts from the group’s mean for each slice were calculated, and the mean and standard deviation of these metrics across all slices and all individuals are reported.

### Development of an automated crypt counting workflow

The automated crypt counting workflow (AW) developed in this work (**Fig. 2**) consists of the following steps.

**Figure 2.**
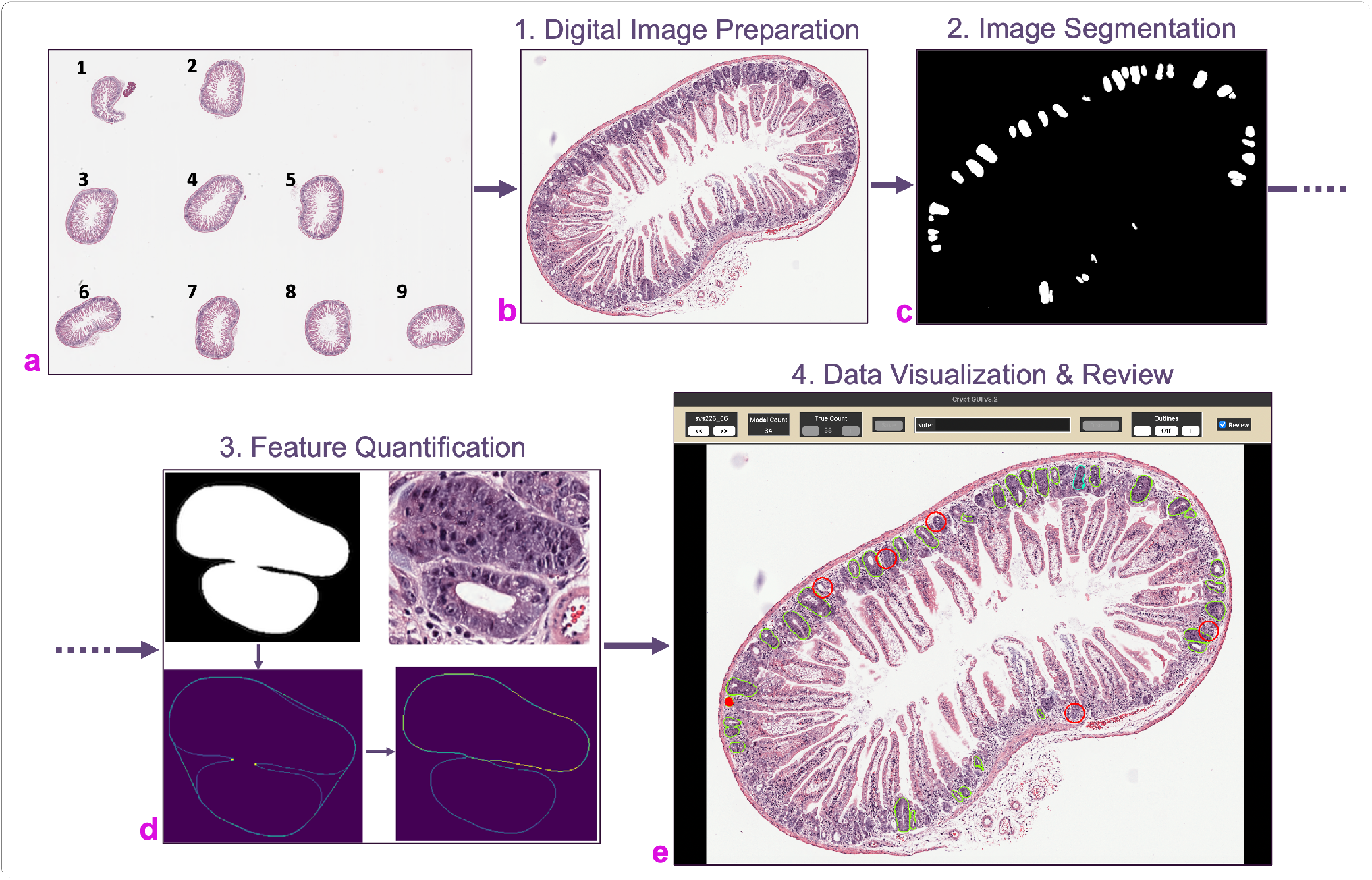
Automated crypt counting workflow. After irradiation, samples are obtained, prepared, and subjected to microscopic imaging, yielding **(a)** a digital whole-slide image (WSI) of all 9 jejunal slices from that mouse. Next, the automated workflow take places in 4 steps: (1) Digital Image Preparation, in which each slice of the WSI is automatically cropped and normalized for resolution and color to the stain intensity, yielding a **(b)** PNG image of each slice. (2) Image Segmentation: each slice image is input into a segmentation model, which yields a **(c)** binary mask of crypt locations for that slice image. (3) Feature Quantification: crypts are identified and enumerated in the segmentation mask. **(d)** Each contiguous image segment (top left) from the mask is analyzed. If any patterns are detected (bottom left) that indicate that a segment comprises two adjacent crypts, the segment is delineated (bottom right), and the crypt count is adjusted accordingly. (Top right) the corresponding crypts in the actual slice image. (4) Data Visualization & Review: the **(e)** graphical user interface is used to examine the automated crypt counts and segmentations of each slice, discard any faulty images, and edit the automated counts and segmentations (green outlines). Corrections made by the user are shown in red.

#### 1) Digital Image Preparation

A Python script was used to load each WSI **(Fig. 2a)**, automatically detect the borders of the 9 jejunal slices, then crop and save 9 full-color, full-resolution images, one of each slice, in Portable Network Graphics (PNG) file format **(Fig. 2b)**. Each slice image is normalized for resolution to account for potential variations in the microscope zoom setting and for color to account for potential variations in H&E stain intensity as described elsewhere [2].

#### 2) Image Segmentation

A segmentation model was constructed with nnU-Net to determine crypt pixels in the PNG images of jejunal slices. To train the segmentation model, nnU-Net requires input images with corresponding segmentation masks, which share the same pixel orientation as the original image, but instead contain an integer for each pixel denoting which structure that pixel belongs to. In our case, binary segmentation was used to indicate crypt or no crypt.

Dataset B (**Table 1**) was chosen for development of the segmentation model. To maximize the diversity of the images in this dataset, slices were chosen from various irradiated mice over 5 different trials and encompassed a large range of numbers of crypts per slice. Individual slices from each mouse were considered independently from each other, and often only a few slices per mouse were selected for this dataset. For each image, 3 individuals with expertise in MW attained consensus regarding the crypt locations, as described in the criteria outlined in the MW protocol, to create a binary mask of the crypt locations by using 3D Slicer software [15]. Of the resultant 59 images and corresponding segmentation masks of Dataset B, 54 images served as the training dataset and were used to train a segmentation model with nnU-Net on GPU nodes of a high-performance computing cluster; the other 5 images were reserved for testing only. nnU-Net implements a 5-fold cross-validation training method, in which the training dataset is separated into 5 subsets, and 5 separate models are trained, each on 4/5 of the subsets using the remaining subset for validation, before being combined into a single model.

Within the automated crypt counting workflow, each slice image produced by the digital image preparation step is input to the segmentation model, which outputs a binary mask image indicating the pixels that make up the crypts in each slice image **(Fig. 2c)**. These predictions are performed on GPU nodes of a high-performance computing cluster.

#### 3) Feature Quantification

Crypt counts of each segmentation mask produced by the segmentation model are extracted via an algorithm written in Python. Different features that can be extracted at this stage include the number of crypts, the area of each crypt, and the area or circumference of the jejunal slice. To extract these data, the borders of each individual crypt from the binary mask are first determined by using a border-following-algorithm described elsewhere [16]. Since the segmentation model does not distinguish between neighboring crypts, it cannot be assumed that each connected pixel area is a single crypt. Rather, it is possible that a single area contains more than one adjacent (touching) crypts. Therefore, each segment is further analyzed with an adjacent-crypt discrimination algorithm (described in the Supplementary Materials) to predict how many crypts it comprises, and to approximate where the borders of those crypts lie **(Fig. 2d)**. Also, any segment smaller than 2000 pixels (500 µm^2^) is assumed to be a false positive and is ignored. The pixel coordinates of the borders of each detected crypt, as well as any other resultant data such as the area of each crypt and the total number of crypts, are saved to a file.

#### 4) Data Visualization and Review

A graphical user interface (GUI) was developed in Python that allows users to view an overlay of the borders of each crypt, as determined by the segmentation model and the adjacent-crypt discrimination algorithm, on the original slice image **(Fig. 2e)**. This overlay makes it easy to detect any errors introduced by the image segmentation and adjacent-crypt discrimination steps. The user can then use the interface to mark any problems, correct the total crypt count, and automatically save these corrections to an Excel spreadsheet. At this step, any incomplete or distorted slice, resulting from errors in either the physical placement of the slice on the slide or in imaging, can be identified and removed from the analysis.

### Evaluation of the automated workflow in comparison with the manual workflow

The AW was evaluated in 3 ways for comparison with the MW. First, the crypt enumeration algorithm of the AW was evaluated by applying the AW to the training (n = 54) and test (n = 5) subsets of Dataset B. The training subset was used to evaluate accuracy in an ideal scenario where the training data perfectly represented future applications of the crypt model, whereas a separate test dataset assessed generalizability to new data. Second, the accuracy of the AW was assessed by directly comparing un-edited AW counts with MW counts across the 3 expert individuals on Dataset A2. In this analysis, we were interested not only in AW and MW count alignment on additional slices not seen during model training, but also in the difference in time between the two workflows. Third, we investigated the discrepancies between the AW and the MW in detail by applying the AW to Dataset C and allowing an expert to manually review and correct the automated counts via the GUI. This analysis was also done by a different expert on Dataset D. Here, we were interested in how many and what types of corrections the experts thought were necessary, as well as the time spent reviewing each image. The latter metric was achieved by implementing a timer built-in to the GUI, to which the individuals were blinded.

### Evaluation of novel metrics for the microcolony survival assay

We also considered two automated metrics in addition to the standard crypt count: (a) the crypt count normalized to the jejunal slice circumference and (b) the crypt count normalized to the average crypt area of the slice. The former metric was computed by dividing the automated crypt count of a jejunal slice by an automated measurement of that slice’s circumference (described further in the Supplementary Materials) and aimed to account for variations in the jejunal diameter. The latter was computed by dividing the automated crypt count of a jejunal slice by the average area, in pixels, of the crypts on that slice, and aimed to account for variations in crypt sizes. To test for any systematic dependency of the slice circumference on its order along the jejunum, the correlation between slice circumference and slice number of all slices in Dataset A2 was computed. To evaluate which metric more closely correlated with radiation damage, we compared the percent standard deviation from the mean across the 5 mice in each treatment group of Dataset A2 for each of the following metrics: the manual counts of each of the 3 experts, the automated crypt count, the slice-circumference-normalized crypt count, and the crypt-area-normalized crypt count. To make them quantitatively comparable, each new metric was multiplied by a constant scaling factor, arbitrarily chosen to yield the closest absolute values to the automated counts. This scaling factor made no difference to the quantitative evaluation, and served only to better visually represent the data.

## Results

### Evaluation of the Manual Crypt Counting Workflow

Trained individuals’ crypt counts across the jejunal slice images of Dataset A deviated from their own mean by 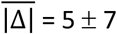, and from the group mean by 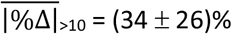 **(Fig. 3)**. The disproportionately high 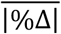of slices with low (<10) crypt counts is evident in **Figure 3**.

**Figure 3.**
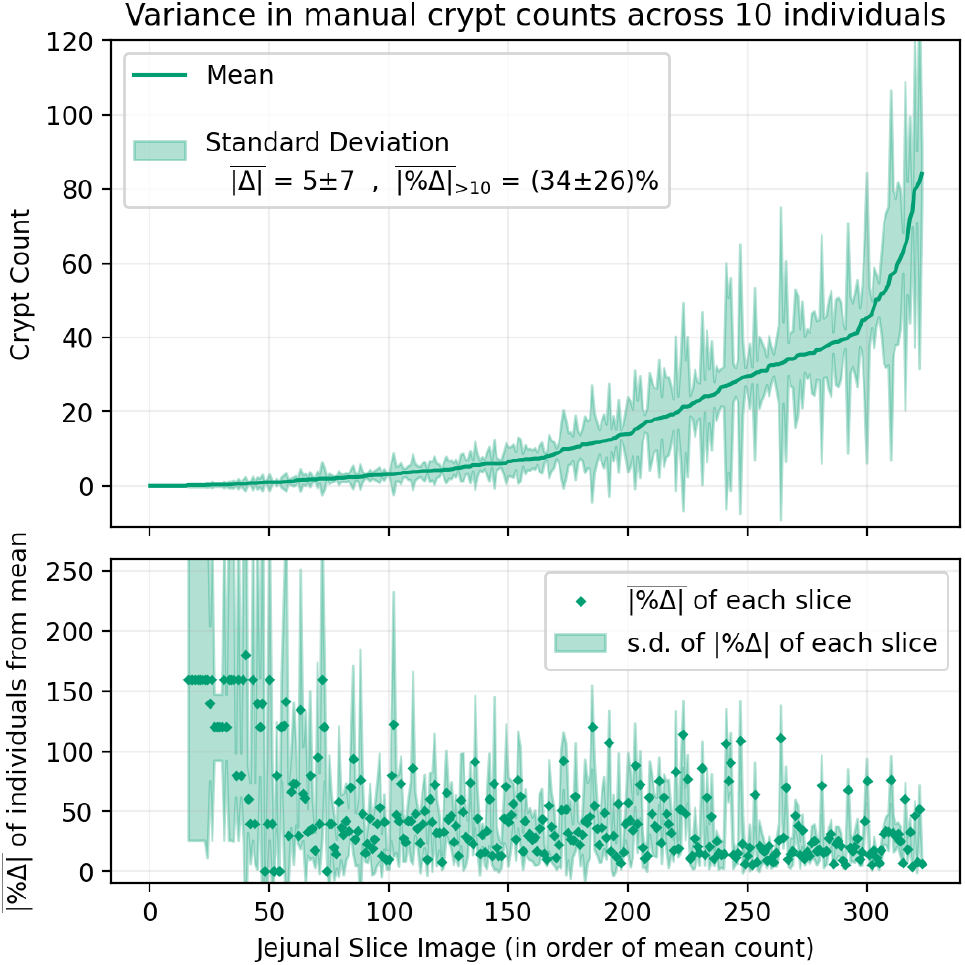
(top) Means and standard deviations between 10 individuals counting crypts in jejunal slice images in Database A. (bottom) For each slice image, the mean and standard deviation of the absolute percent deviations of all individuals from their mean are shown. Note that the top and bottom plots of this figure share an x-axis.

### Development of an Automated Crypt Counting Workflow

An automated workflow was developed that consisted of the following steps: (1) digital image preparation, (2) image segmentation, (3) feature quantification, and (4) data visualization and review.

During the training of the segmentation model constructed by using nnU-Net for crypt segmentation, the mean Dice coefficient across the 5 training folds was 0.88 ± 0.01. The training took ∼42 days of CPU time and used a maximum of 9 GB of RAM. The mean size of each training image was 23 ± 5 MB. Running predictions of the segmentation model onto its own 54 training images from Dataset B yielded a mean Dice value of 0.99 ± 0.00. Running predictions of the model onto the 5 testing images of Dataset B yielded a mean Dice value of 0.89 ± 0.02. A critical component of the feature quantification step was the adjacent-crypt discrimination algorithm, which aimed to identify any groups of crypts with shared adjacent borders within the segmentation model output and delineate and count the individual crypts within each group. The true crypt count for each jejunal slice image in the training dataset versus the automated workflow count, either with or without the adjacent-crypt discrimination step, is shown in **Figure 4**. Without discrimination, the automated workflow tends to undercount [i.e., 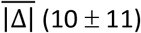 and 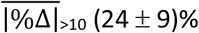], worsening substantially for slices with higher total crypt counts. With discrimination, the automated workflow did not undercount or overcount and remained closer to the true count 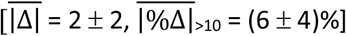.

**Figure 4.**
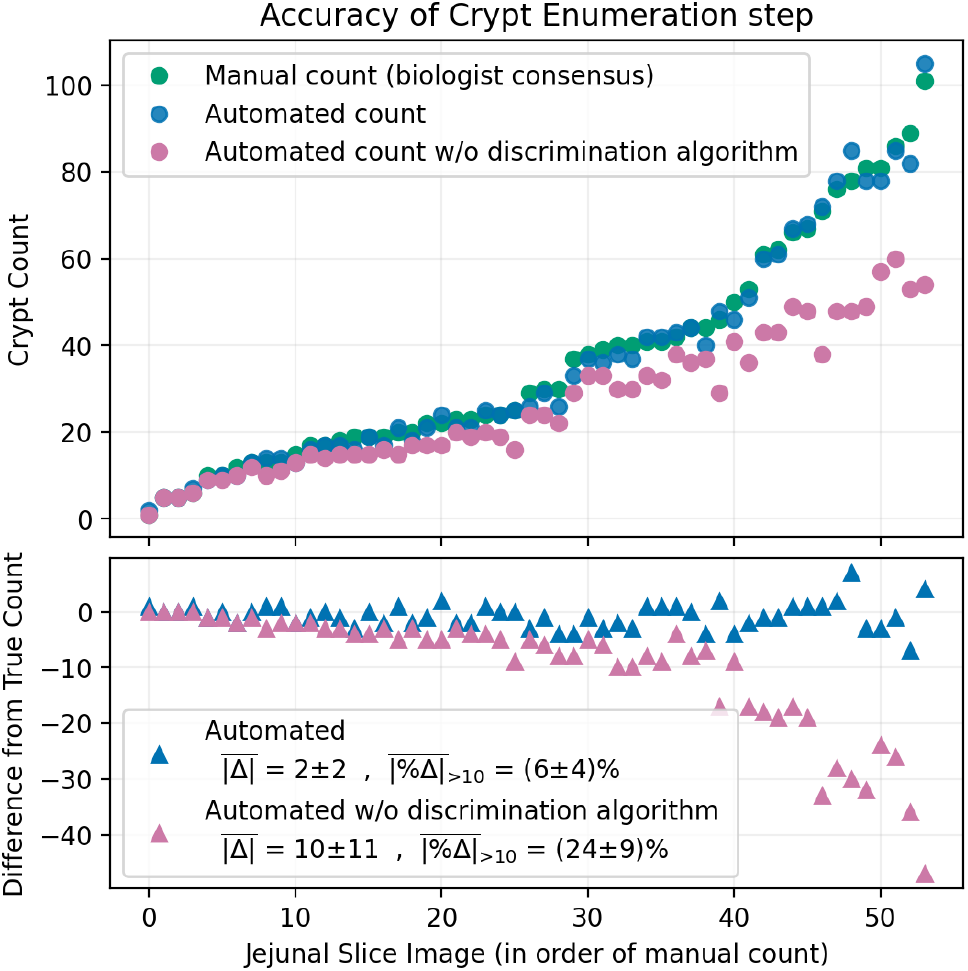
Manual workflow crypt counts versus the automated workflow’s crypt counts, both with and without the adjacent-crypt discrimination algorithm, of the training images from Dataset B. Manual counts were derived by consensus of 3 experts who manually segmented the training data. Note that the top and bottom plots of this figure share an x-axis.

The entire automated workflow took a few hours to complete for a typical dataset of 60 WSI, necessitating only a few minutes of active time by the user. Most of the time was taken by the digital image preparation and image segmentation steps, as well as the time to transfer files to and from the high-performance computing cluster. The feature quantification step, in contrast, took only a few minutes per dataset. Computer time while using the GUI was negligible, loading images and their associated data within a couple of seconds.

### Comparing the automated workflow with the manual counting workflow

The variation in the manual counts of the 3 expert counters of Dataset A2 deviated from their mean by 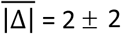 and by 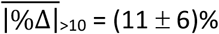 **(Fig. 5)**, which was lower than the deviation observed across all 5–10 individuals who manually counted Dataset A. Relative to the mean manual counts of this subset of Dataset A, the automated counts deviated by 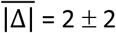 and 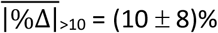. The largest discrepancies between the two counting workflows tended to occur on slice images with large variance among the 3 individuals’ counts, as indicated by the propagated uncertainty in the difference calculation [the shaded areas on **Fig. 5 (bottom)**], which only rarely did not include 0. The automated counts deviated by 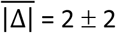 and 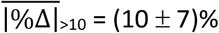 from the first expert; 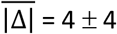 and 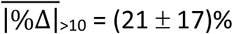 from the second expert;and 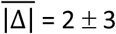 and 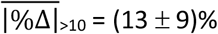 from the third expert.

**Figure 5.**
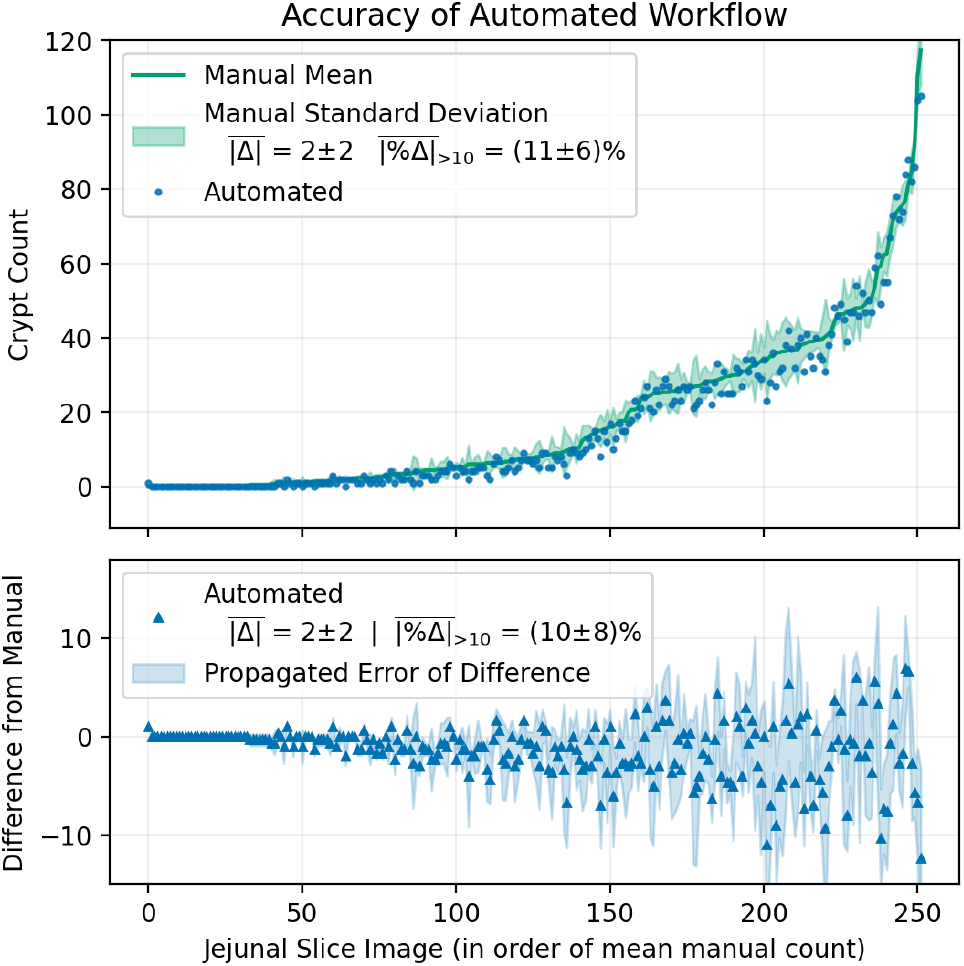
Comparison of the automated workflow’s crypt counts to the mean of the 3 expert individuals’ crypt counts of a subset of Dataset A. (bottom) The differences between the automated counts and mean manual counts. Note that the top and bottom plots of this figure share an x-axis.

### Evaluation of the automated workflow by manual correction

Automated counts of jejunal slice images from Dataset C were reviewed and corrected by an expert counter by using the GUI built into the automated workflow. The same was done by a different expert for Dataset D. In Dataset C, the automated counts differed from the manual corrections by 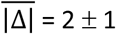 and 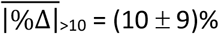, tending to overcount for slices with more than 10 crypts **(Fig. 6a)**. In Dataset D, the automated counts differed from the manual corrections by 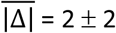 and 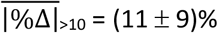, tending to undercount on slices with <40 crypts, and overcount on slices with >40 crypts. In Dataset C, the individual spent an average of (52 ± 21) s reviewing each slice image, as indicated by a timer built in to the GUI. Retrospective analysis of the review times identified instances in which the individual was interrupted in the counting workflow, resulting in outliers not indicative of the time spent reviewing. Thus, images with review times >100s, which made up 22% of all images in the dataset, were removed from the analysis. In Dataset D, the individual was measured to have spent an average of (29 ± 16) s reviewing each slice image, with no outliers requiring removal.

**Figure 6.**
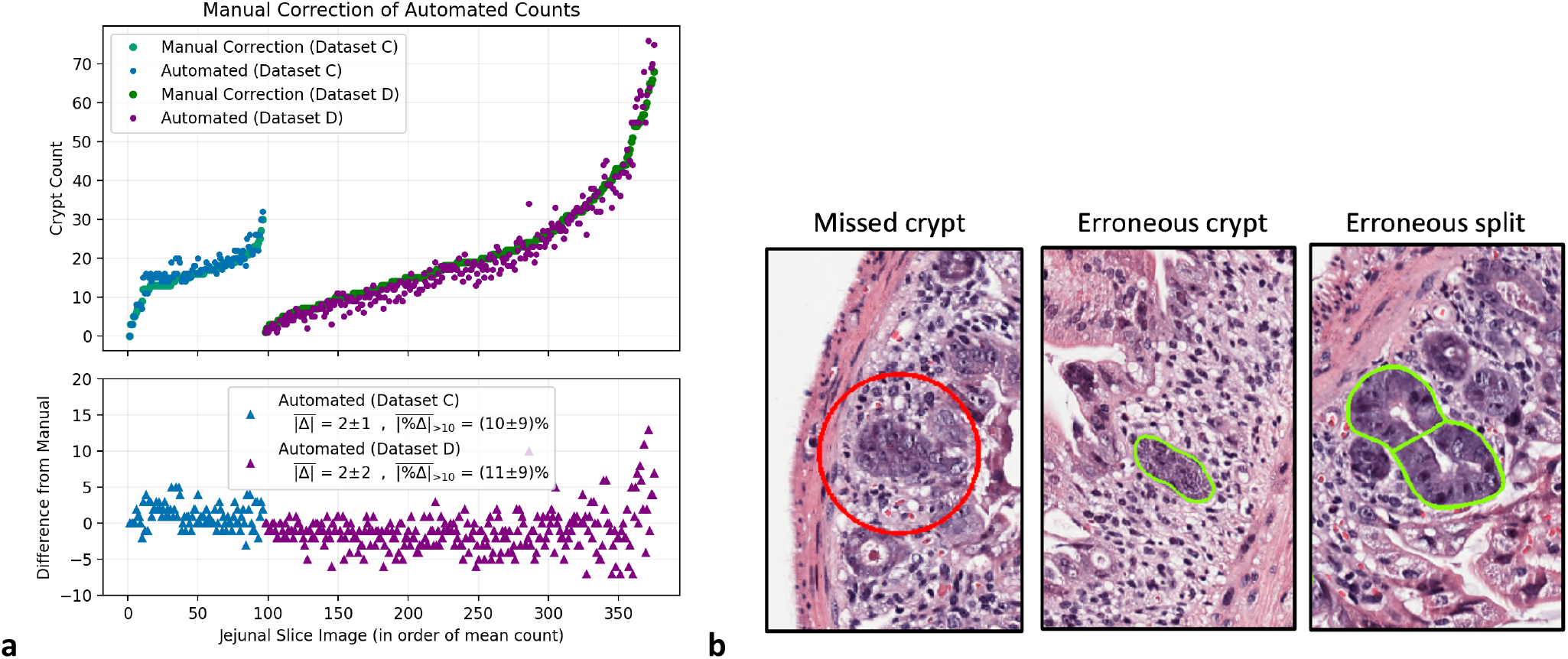
**(a)** (top) Comparison of the automated workflow’s crypt counts of Dataset C and Dataset D to the manual corrections performed by an expert individual on either dataset. (bottom) The differences between the automated and manually corrected counts. Note that the top and bottom plots of this subfigure share an x-axis. **(b)** Examples of the various error types identified during review: missed crypt, erroneous crypt, and erroneous split.

An error type analysis was carried out by both reviewers. Of the 100 total slice images in Dataset C, 4 were removed from the analysis, 3 because the slice was broken or torn, and 1 for an issue with the microscope image. Of the 306 total slice images in Dataset D, 28 were removed from the analysis, 24 because the slice was broken or torn, 2 for issues with the microscope image, and 2 for poor staining. The errors present in the remaining images were classified as either missed crypt, erroneous crypt, or erroneous split **(Fig. 6b)**. ‘Missed crypt’ refers to crypts that the reviewer deemed to fit the criteria to count but were not identified by the AW; ‘erroneous crypt’ refers to crypts that the reviewer did not deem to fit the criteria to count but were identified by the AW; and ‘erroneous split’ refers to crypts that were correctly identified by the AW, but where the adjacent-crypt discrimination algorithm erroneously split the crypt into two. The frequencies of the various error types for each dataset are shown in **Table 2**.

**Table 2.** Percentage of total crypts with given error types, as identified through manual review of Datasets C and D by exper.

| Dataset | Total No. of Crypts | No Errors | Missing Crypt | Erroneous Crypt | Erroneous Split |
| --- | --- | --- | --- | --- | --- |
| C | 1,513 | 87% | 3% | 2% | 8% |
| D | 5,961 | 81% | 12% | 1% | 6% |

### Evaluation of novel metrics for the microcolony survival assay

A positive Pearson correlation coefficient of *r*=0.22 (two-sided *p* value 0.0005) was found between slice circumference and slice number along the jejunum for all slices of Dataset A2 (**Fig. 7a**). The average percent standard deviation of each slice number’s circumferences was 9%. The mean across all 9 jejunal slices of each mouse in each treatment modality of Dataset A2 for each metric (manual crypt count of each expert, automated crypt count, slice-circumference-normalized automated crypt count, and crypt-area-normalized automated crypt count) is plotted with circular marker points in **Fig. 7b (top)**. The latter two metrics were scaled with arbitrary scaling factors of 11,000 and 10,000, respectively. The mean and standard deviation across the mice in each treatment modality for each metric is depicted by a bar graph and associated error bars. The average relative standard deviation across treatment modalities for each metric is compared in **Fig. 7b (bottom)**.

**Figure 7.**
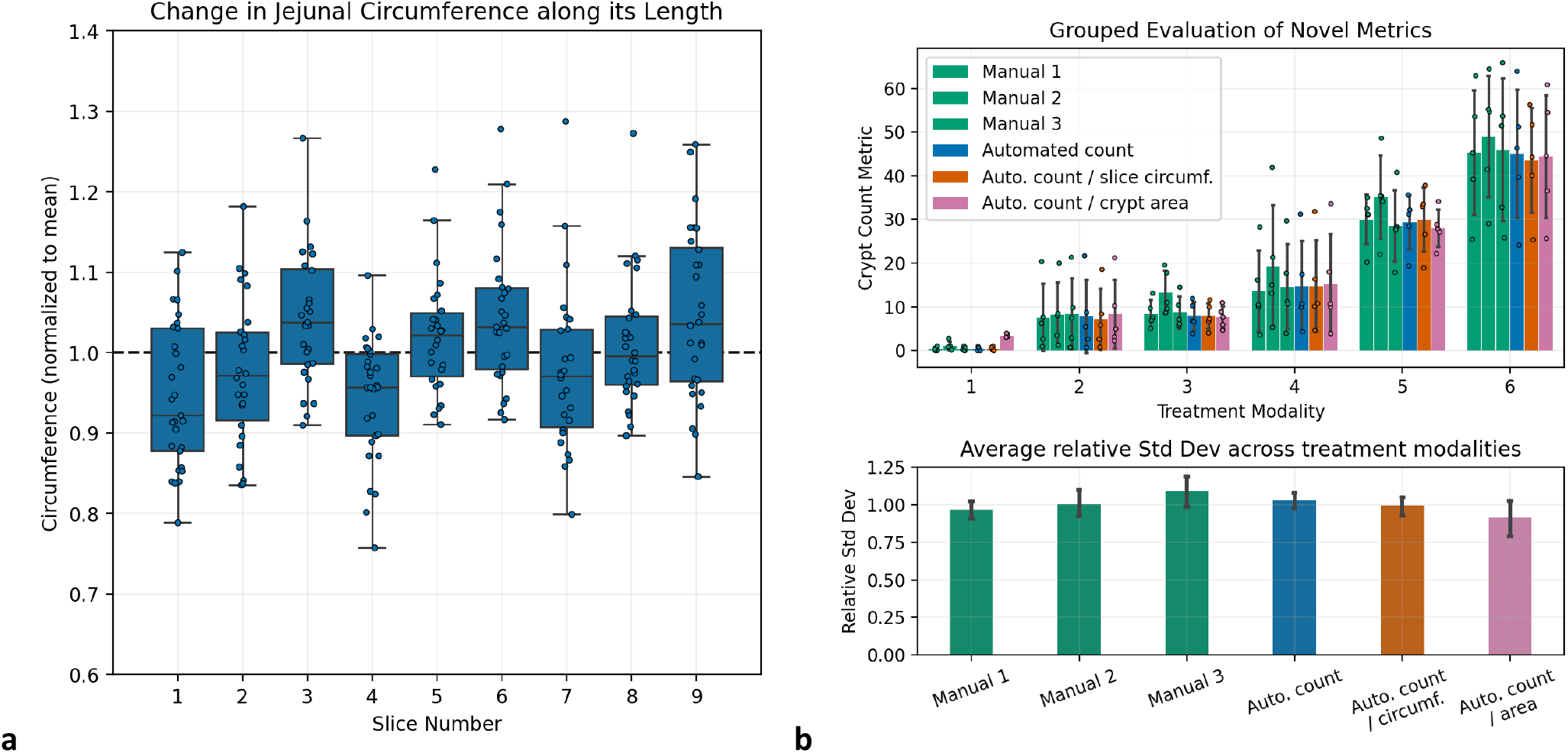
**(a)** Boxplot of circumferences of all jejunal slices of Dataset A2 as a function of slice number (along the jejunum), normalized to the mean circumference. **(b)** (top) Comparison of each of the 3 experts’ manual counts to the automated workflow’s crypt counts, the slice-circumference-normalized automated crypt count, and the average-crypt-area-normalized automated crypt count of Dataset A2, grouped by treatment modality. Individual data points show the mean crypt count across all 9 jejunal slices of each of the 5 mice per treatment modality. Colored bars and associated errors show the mean and standard deviation across the 5 mice. (bottom) The average relative standard deviation of each count type across all treatment modalities except for modality 1.

Treatment modality 1 is excluded from this evaluation because the near-zero absolute counts disproportionately skewed the relative standard deviations. No metric had a consistently lower percent standard deviation than the other metrics across the treatment modalities.

## Discussion

The microcolony survival assay remains an important assay for radiation biology research but is limited by the tedious and time-consuming nature of manual analysis and the confounding effect of intra– and inter-observer variability. The latter issue is evident in the high variance 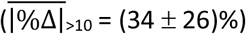 in crypt counts between trained individuals using the manual workflow. In fact, this variance may be larger than the typical differences in crypt counts observed between treatment cohorts in relevant radiobiological studies [17-21]. Because the variance across the 3 expert counters was lower 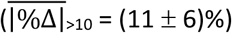, we can deduce that these inconsistencies may be improved by further training but that inherent differences due to subjective evaluation remain. These results strongly underscore the need for a fast and robust workflow.

To address these shortcomings, we automated the microcolony survival assay in its entirety, mirroring how it would be performed traditionally. We focused explicitly on providing full transparency over the automation processes and enabling human intervention to facilitate laboratory adoption and trust in the automated assay. Our AW consisted of 4 steps: (1) digital image preparation, (2) image segmentation, (3) feature quantification, and (4) data visualization and review.

The digital image preparation step standardized the resolution and stain color intensity between different images to improve the generalizability of the AW to variations in imaging and staining protocols. Also, separating the WSI into images of its individual slices allowed subsequent analysis to be randomized to reduce the potential bias present in the MW from examining all slices of each mouse together. The image segmentation step, the computer identification of the structures of interest (crypts), is perhaps the most fundamental component of the AW yet was the simplest to develop. Typically, training an AI segmentation model requires relevant expertise to select the optimal model architecture and train parameters for the dataset at hand. Using nnU-Net obviated these requirements, as it is a self-configuring framework that automatically chooses and implements the optimal parameters for the entire model training pipeline, including image pre-processing and U-Net deep learning network architecture selection [12]. The resultant segmentation model yielded highly accurate predictions on the test dataset (mean Dice coefficient 0.89). However, some overfitting was present, with the model obtaining a mean Dice coefficient of 0.99 on the training data and plateauing in the validation set performance at ∼10% of the total training iterations across all 5 cross-validation folds. Developing the feature quantification step did require specific image processing expertise, most notably for the adjacent-crypt discrimination algorithm. Without this algorithm, the AW would fail to recognize adjacent crypts as individual entities and thus undercount, especially in slices with higher crowding of crypts due to high crypt counts. The crypt enumeration algorithm performed well, though imperfectly, on segmentations of the training dataset, deviating from the true count by 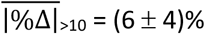. The inaccuracies were caused by the erroneous splitting of oddly shaped crypts and the erroneous non-splitting of some adjacently touching crypts. The GUI facilitated the review and correction of the automated data, and made human error in the recording of data virtually impossible. The minimum size threshold of 2000 pixels (500 µm^2^) for a segment to be counted as a crypt was chosen based on the 10 cell minimum outlined in the criteria; the sizes of the smallest cells observed in viable crypts were ∼50 µm^2^ each.

Comparing the AW to the MW was complicated by the inherent user-dependent variance of the MW. Thus, the AW was first compared to the mean of 3 experts’ MW counts, deviating by 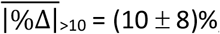, which was less than the mean deviation of the 3 experts from their own mean 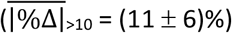. This indicates that the AW was at least as accurate as any one expert, and more accurate than the average trained individual, in computing the MW mean counts. Expert review of the automated counts on separate datasets revealed similar deviations of the AW from the two experts’ corrections 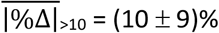 and 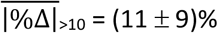, indicating that the magnitude of this deviation is robust across various datasets. The tendency of different experts to either overcount or undercount relative to the AW may be indicative of inter-observer subjective variability. In fact, the subjectivity of even expert analysis is illuminated by the different error types each expert identified.

Although the experts found similar rates of erroneous splits and erroneous crypts, which were typically easy to identify, they differed substantially in their identified rates of missed crypts. This task was arguably the most subjective, with crypt-defining criteria including “being basophilic, having a clear structure, and being located on the circumference of the slice.” Also worth noting is that the deviation between the automated and manually corrected counts of each slice occasionally underestimated the actual number of crypt errors identified in the slice by the reviewer, because a slice could coincidentally have both overcount errors (e.g. erroneous split) and undercount errors (e.g. missed crypt), resulting in a net balance of errors.

We evaluated two novel metrics, the slice-circumference-normalized crypt count and the average-crypt-area-normalized crypt count, in an attempt to identify a normalization metric that could improve the microcolony survival assay in terms of correlating histopathological features to radiation damage. Because crypt count could be expected to increase linearly with jejunal circumference and independently of radiation damage, normalizing to the circumference seemed reasonable. In our study, the circumferences of slices across Dataset A2 varied consistently by 9%. Thus, normalization could be expected to noticeably affect the results.

Normalization to crypt area has been recommended by prior studies, because the average crypt size has been shown to increase after radiation damage and the total number of cells per crypt to increase by 30%–50% after radiation [20]. Techniques involving radioactive staining of proliferating cells have been described that can account for variations in crypt size [18, 22]. To evaluate which normalized metric more closely correlated with radiation damage, we made the following assumptions: (1) ‘Biological variance’ is inherent in the phenotype of radiation damage observed in mice receiving the same treatment; (2) each metric is associated with ‘metric variance’, which is independent from the ‘biological variance’. Therefore, the metric most closely correlated to radiation damage will have the lowest ‘metric variance’, resulting in total variance that primarily reflects biological variance alone. In this study, no metric demonstrated a consistently lower total variance than the other metrics; thus, this evaluation did not indicate that any metric was more or less correlated with radiation damage than the standard crypt count. Normalization was thus not beneficial in this context. However, the method used to compare the metrics should be scrutinized, because the relative impacts of the assumed ‘metric variance’ and ‘biological’ variance are unknown and may render the method ineffective. A better method would be to directly examine the correlation between the metric and a known metric of radiation damage, such as radiation dose, as was originally done by Withers and Elkind to establish the assay [1].

The application of computer automation to this assay has been explored previously. Fu et al. correlated jejunal slice images of irradiated mice to the doses they were irradiated with and trained a deep learning model to predict the delivered dose based on any given jejunal slice image [11]. They observed that their deep learning model led to more precise dose estimations for some of the tested dose points compared with the manual crypt counting method, albeit at the expense of a less accurate dose estimation for other tested dose points.

Although partially successful in terms of implementation, the approach chosen by this group presents some limitations. First, as mentioned in their report, by correlating each image directly to the delivered dose, and not to a biological feature indicative of damage (e.g. crypt count), the application of the automated assay must remain limited to the parameter domain defined by their training data. For example, if a new radiation modality (or a non-radiation treatment modality) were to be compared outside of the ones used in their training data, it is unclear how that would affect the appearance of the jejunal slices in the resultant images and how these changes would be interpreted by the deep learning model. Because the model features indicative of greater or lesser dose burden are unknown, they may not be correlated with biological features independent of the radiation modality. Thus, although such a model may be useful in specific applications, it cannot fully replace the widely generalizable standard microcolony survival assay. Second, again because of the lack of a biological feature on which the automated assay is founded, the output of the automated assay cannot be reviewed by human users. Instead, the model must be fully trusted when applied to new data, because no opportunity exists for a redundant check of the model output. Again, any variation in sample preparation (e.g., staining or slice placement on glass slides), in the slice samples themselves (e.g., differences between mice), or in the effect of the new treatment on the jejunal slice could cause unpredictable and erroneous variation in the model output. These issues would also complicate the adoption of this automated assay in other laboratories, which would have to create their own standard curves as well as potentially retrain a new model if their sample preparation standards differ enough. Liu et al. developed a semi-automated assay to assess intestinal radiation toxicity on histopathologically stained samples of mouse jejunum, not using the microcolony survival assay but rather by quantifying the amount of proliferation, villi length, and DNA damage.

In the current study, although the AW performed as well as or better than the average expert in predicting the manual mean counts, aspects of the AW could be improved. The most obvious area for potential improvement is in the feature quantification step, specifically the adjacent-crypt discrimination algorithm. Both human reviewers identified erroneous splits responsible for a significant fraction of identified errors. The algorithm would often split a single crypt into two (as exemplified in **Fig. 5b**) if the crypt’s outline had concavities. Sometimes, it would coincidentally split multiple crypts into the correct number of constituent crypts, but not along the actual borders of the adjacent crypts. Less often, the algorithm would fail entirely to split multiple crypts into their constituents. These issues could be addressed by further refining the rules followed by the algorithm, for example by considering that adjacent crypt borders are typically parallel to hypothetical radial lines from the center of the slice to its circumference. Alternatively, the overfitting present in the image segmentation model could likely be improved by fine-tuning the model architecture or limiting the number of training iterations. This could result in both less missed crypts and less erroneous crypts, and could also improve the adjacent-crypt discrimination algorithm if crypt borders were more accurately segmented.

However, the most promising potential improvement to the image segmentation step would be to implement an instance segmentation model, as opposed to the semantic segmentation model produced by nnU-Net.

Whereas semantic segmentation only classifies each pixel (in our case the classification was binary—either crypt or non-crypt), instance segmentation also distinguishes between multiple instances of the same class. An instance segmentation model could recognize adjacent crypts as separate entities, and the subsequent crypt enumeration step would become trivial because no discrimination would be needed. Nonetheless, we emphasize the high accuracy of image segmentation in spite of these limitations. The rest of the AW, the digital image preparation and data visualization steps, did not seem to require improvements.

In this report, we describe developing a workflow to automate the microcolony survival assay. In addition to saving time and tedious manual labor, the AW performed the assay more accurately than the average trained manual counter, and just as accurately as the average expert manual counter. Avenues for potential improvement of the AW were identified, most notably the implementation of an instance segmentation model instead of the current semantic segmentation model. AW deployment requires minimal effort and financial investment, facilitating its adoption by reseachers and perhaps establishment of an inter-institutional standard. A similar automated approach could be applied to other histopathological assays.

## Acknowledgements

We thank Christine F. Wogan, MS, ELS, of MD Anderson’s Division of Radiation Oncology, for editorial contributions to this article.

