## Supplementary Materials for "Modernizing histopathological analysis: a fully automated workflow for the digital image analysis of the intestinal microcolony survival assay"

**Adjacent-crypt discrimination algorithm**

The adjacent crypt discrimination algorithm analyzed segments from the semantic segmentation model and attempted to identify the number of crypts comprising each segment and approximate their respective borders. For each binary segment, e.g. **Figure 2d (top-left)**, the algorithm took the following steps:

1. Compute the border (contour) of the segment with cv2.findContours
2. Compute the convex hull of the contour with cv2.convexHull (see outer blue outline of **Figure 2d (bottom-left)**)
3. Compute any concave defects from the convex hull with cv2.convexityDefects (see yellow points of **Figure 2d (bottom-left)**)
4. Depending on the number of defects, their distance from the convex hull, and their orientation relative to the shape of the segment, decide to split the segment by a straight line between given defects
5. If a decision to split was reached, redefine the contour as two separate contours sharing a straight-line border along the split **Figure 2d (bottom-right)**).
6. Iteratively repeat this process on the segment until no more splits are required

**Automated jejunal slice circumference measurement**

To automatically compute the circumference of each jejunal slice (e.g. **Figure S1**), an algorithm took the following steps:

1. Binarize the image and apply an intensity threshold to identify the pixels comprising tissue
2. Find the border (contour) of the largest contiguous segment of the tissue mask with cv2.findContours
3. Find that contour's convex hull with cv2.convexHull
4. Compute the length, in pixels, of the convex hull

a) 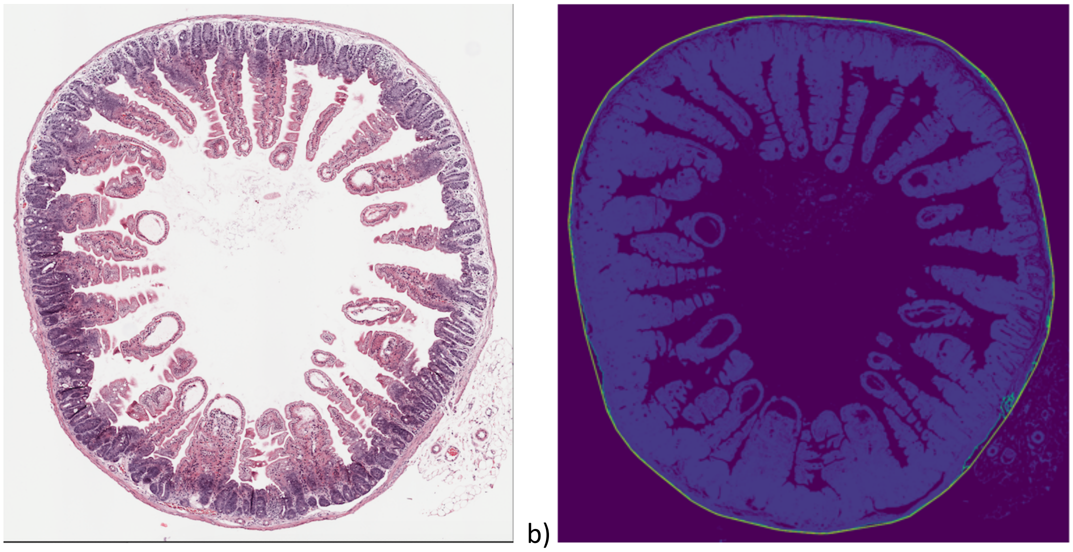 b)

**Figure S1. (a)** Example of a jejunal slice image. **(b)** The tissue mask (blue shaded regions), the tissue outline (bright green outline, barely visible in bottom right portion of the slice), and the convex hull (bright yellow outline). The slice circumference is given by the length of the convex hull.
